# Hypoxia-mediated suppression of pyruvate carboxylase drives tumor microenvironment immunosuppression

**DOI:** 10.1101/2022.04.06.487050

**Authors:** Michael F. Coleman, Eylem Kulkoyluoglu Cotul, Alexander J. Pfeil, Emily N. Devericks, Hao Chen, Violet A. Kiesel, Muhammad H. Safdar, Dorothy Teegarden, Stephen D. Hursting, Michael K. Wendt

## Abstract

Metabolic reprogramming and immune evasion are established hallmarks of the tumor microenvironment (TME). Growing evidence supports tumor metabolic dysregulation as an important mediator of tumor immune evasion. High TME levels of lactate potently suppress antitumor immunity. Pyruvate carboxylase (PC), the enzyme responsible for the anaplerotic conversion of pyruvate to oxaloacetate, is essential for lung metastasis in breast cancer. Moreover, PC may be dispensable in some cells within the TME, and loss of PC expression is associated with immunosuppression. Here we test whether PC suppression alters tumor metabolism and immunosuppression. Using multiple animal models of breast cancer, we identify a dimorphic role for PC expression in mammary cancer cells. Specifically, PC supports metastatic colonization of the lungs, while suppression of PC promotes primary tumor growth and suppresses histological and transcriptomic markers of antitumor immunity. We demonstrate that PC is potently suppressed by hypoxia, and that PC is frequently suppressed in solid tumors, particularly those with higher levels of hypoxia. Using metabolomics, high-resolution respirometry, and extracellular flux analysis, we show that PC-suppressed cells produce more lactate and undergo less oxidative phosphorylation than controls. Finally, we identify lactate metabolism as a targetable dependency of PC-suppressed cells, which is sufficient to restore T cell populations to the TME of PC-suppressed tumors. Taken together, these data demonstrate that elevated lactate following PC suppression by hypoxia may be a key mechanism through which primary tumors limit antitumor immunity. Thus, these data highlight that PC-directed tumor metabolism is a nexus of tumor progression and antitumor immunity.

## INTRODUCTION

Reprogramming metabolism and escaping immune destruction are hallmarks of cancer (1). The interplay of cancer cells and immune cells in the tumor microenvironment (TME) is a critical determinant of cancer progression and treatment efficacy (2,3), and is mediated in part through metabolic interactions (4-6). Rapid cell proliferation and vasculature dysfunction frequently result in solid tumor hypoxia and subsequently reprogramming of central carbon metabolism (7). Such hypoxia promotes local and metastatic spread via numerous mechanisms including metabolic and transcriptional reprogramming of the TME (7). Tumoral hypoxia also alters the TME immune landscape by dampening antitumor immune responses while supporting protumor inflammatory signaling (8).

The rapid metabolism of cancer cells can lead to a TME with low concentrations of necessary anabolic nutrients such as glucose and glutamine (6). Such TME remodeling poses a hostile metabolic environment for activated cytotoxic T cells, key mediators of antitumor immunity, which compete with cancer cells for glucose and glutamine (6). In contrast, T regulatory (Treg) cells rely more heavily on fatty acid oxidation, thus, enabling the accumulation of Treg cells in the metabolically hostile TME (9). Lactate, produced by cancer cells at high rates, promotes both formation and activity of Treg cells to limit antitumor immunity (10) and restrict cytotoxic activity of CD8^+^ T cells (6,11). Indeed, high lactate levels in the TME regulate many cell types within the TME, including cancer cells, T cells, NK cells, tumor associated macrophages (TAMs) and dendritic cells to limit antitumor immunity (12-16). Lactate secretion from tumor cells is facilitated via monocarboxylate transporters (MCTs) and is accompanied by the export of H^+^ ions acidifying the TME to drive further immunosuppression (17,18). Therefore, the expression and activity of these transporters play a pivotal role in immune cell function in the TME and are emerging as important drug targets for the treatment of solid tumors (16,19-21).

Pyruvate carboxylase (PC) converts pyruvate into oxaloacetate (OAA) to replenish the tricarboxylic acid (TCA) cycle (22). PC is essential for the development of pulmonary metastasis, with higher expression in pulmonary metastases compared to non-pulmonary tumors (23-25). However, the relationship between PC and primary tumor growth is less well understood. Pharmacological inhibition of PC, using both immune intact and immunocompromised animals, reduced primary tumor size (23). Conversely, we have found that shRNA-mediated suppression of PC in mammary cancer cells does not alter primary tumor size (24). Similarly, there is no association between PC expression and breast tumor size in METABRIC data (24). Intriguingly PC is suppressed in TAMs from hypoxic tumors, but hypoxia alone does not suppress PC in macrophages (26). Restoration of PC in TAMs promotes T cell-dependent antitumor immunity to limit tumor growth (26).

Thus, while PC clearly supports the pulmonary metastasis in animal models of breast cancer, the relationship between PC, primary tumor growth, and antitumor immunity remains unclear. Hence, we tested whether PC suppression in mammary cancer cells, potentially involving a hypoxia-dependent mechanism, limits antitumor immunity via lactate metabolism.

## MATERIAL AND METHODS

### Cell lines

4T1, E0771, D2.A1, metM-Wnt^lung^, and MDA-MB-231 cells were maintained in Dulbecco’s modified Eagle’s medium (DMEM) supplemented with 10% fetal bovine serum (FBS) and 1% Penicillin-streptomycin. Prior to experiments cells were seeded into media with glucose concentrations were decreased to 5.6 mM. M-Wnt cells were maintained in RPMI containing, 10% FBS, penicillin/streptomycin, 11 mM glucose, and 4 mM glutamine, prior to experiments M-Wnt cells were seeded in human plasma like media (HPLM) containing 10% FBS and 1% penicillin-streptomycin.

PC was suppressed in E0771 and M-Wnt cells by lentiviral transduction for 48 h, using particles generated by transfection of HEK293T cells with psPAX2, pMD2.G, and Pcx-targeting shRNA in pLKO.1 or psi-LVRU6H, respectively. TRC lentiviral mouse *Pcx*-targeting shRNAs (lentiviral pLKO.1 TRC cloning vector) were purchased from GE Dharmacon, (Lafayette, CO). The target shRNA sequences were AAAGGACAAATAGCTGAAGGG (shPC25), TTGACCTCGATGAAGTAGTGC (shPC28), and TTCTCCGAACGTGTCACGT (Scram) and were selected for using puromycin (5 µg/ml). Similarly, Pcx-targeting shRNA were purchased from Genecopoeia (Rockville, MD) in a psi-LVRU6H vector. The sequences were GAGTTGGAAGAGAATTACAC (shPC-B), CCACAACTTCAACAAGCTCT (shPC-C), and GCTTCGCGCCGTAGTCTTA (Scram) and were selected for using hygromycin (100 µg/ml). Doxycycline inducible PC suppression was achieved by transducing a GE Dharmacon SMARTvector encoding Pcx-targeted shRNA (TGCAATCGAAGGCTGCGTA) into M-Wnt cells as described above and selecting with puromycin (5 µg/ml). To generate PC promotor luciferase reporter cells, the PC promotor fragment from pGL3.0-PC-575-luc (27) was subcloned into pGL4.0 expression vector at KPN1 and XHO1 sites, and transfected into cells as indicated with selection puromycin (5 µg/ml). The 4T1, E0771 and MDA-MB-231 cell lines were constructed by stably expressing firefly luciferase under puromycin selection to allow for luminescent imaging of metastatic growth. All cell lines were tested for mycoplasma using R&D Systems MycoProbe Mycoplasma Detection Kit or ATCC Universal Mycoplasma Detection Kit.

### Animal models

All in vivo experiments were conducted in accordance with institutional animal care and use committee protocols approved by the Purdue University or the University of North Carolina at Chapel Hill (Purdue Protocol: 1310000978, UNC at Chapel Hill protocol: 19-227). All mice were provided ad libitum access to food and water. Mice used in E0771 studies were fed D12450J (Research Diets, New Brunswick, NJ). Mice used in M-Wnt studies were fed AIN-93G (Research Diets). Mice used in 4T1-PC-FF studies were fed standard chow diet (Teklad Global 2018S, Envigo, Indianapolis, IN) Mice were monitored by vivarium staff daily for signs of dehydration, pain, or distress.

For pulmonary tumor growth experiments, mice (5/group) were injected via the lateral tail vein with control and PC-suppressed E0771 cells (10^6^/100 µl) and their pulmonary tumor growth was observed with bioluminescence imaging using an Advanced Molecular Imager (AMI) (Spectral Instruments, Tucson, AZ) for 28 days. Mice were subsequently euthanized and the lungs were then removed, and fixed in 10% neutral buffered formalin for 48 hours, and paraffin embedded.

For primary tumor studies, scrambled control and PC-suppressed E0771 (5×10^5^/50 µl) or M-Wnt (5×10^4^/50µl) cells, were orthotopically transplanted in PBS into the fourth mammary fat pad of C57BL6/J mice (n=3-6/group for each experiment). PC promoter-luciferase reporter 4T1 cells (4T1-PC-FF, 2.5×10^4^/50 µl) were orthotopically transplanted into BALB/c mice and grown for 14 days before euthanasia. Body weight was measured weekly and tumor volume (0.5 × length × width^2^) was measured at least twice per week. Primary tumors were weighed; half of each tumor was snap frozen in liquid nitrogen and stored at -80°C; and the other half was formalin fixed and paraffin embedded. Mice bearing doxycycline-inducible shPC were randomized to receive either control or 150 µg/ml doxycycline in their drinking water. In mice undergoing resection of primary tumors, weekly bioluminescence was used to monitor the development of lung metastasis, and lung weight was determined at euthanasia 10 weeks later.

For animals treated with AZD3965, control and PC-suppressed E0771 tumors grew for 12 days and then were randomized to begin daily treatment with AZD3965 by oral gavage (50mg/kg) (MedKoo Biosciences, Morrisville, NC, #1448671-31-5). After 14 days of treatment, mice were euthanized, and tumors processed as above.

#### RNA extraction and qPCR

RNA was isolated from PC-suppressed E0771 and M-Wnt tumors by homogenization in TRIzol, followed by chloroform extraction, and isolation using EZNA HP Total RNA kit (Omega Bio-Tek #R6812, Norcross, GA). For qPCR analyses, cells were grown at a concentration of 2×10^5^ cells/well in corresponding treatment media for 24 hours. The next day, cells were harvested, RNA was isolated (EZNA HP Total RNA kit) and cDNA was synthesized (Thermo Scientific, Verso cDNA Synthesis Kit #AB-1453/B).

Quantitative PCR was performed using Maxima SYBR Green/ROX qPCR Mastermix (Thermo Scientific #K0222) using a Biorad CFX Connect Real Time System (Biorad Laboratories, Inc.). Primer sequences were obtained from the Integrated DNA Technology web site. *Gapdh* or *18S* were used as housekeeping genes to normalize the gene expression level. The relative difference in gene expression level was calculated using the δδcycle threshold method.

### Transcriptomic analysis

Sense-strand cDNA was synthesized, fragmented, labeled, and hybridized onto a Clariom S peg plate (Affymetrix, Santa Clara, CA). The GeneChip® WT PLUS Reagent Kit (Affymetrix) was used to prepare the samples. Labeled cDNA was hybridized to the plate using GeneTitan Hybridization Wash and Stain Kit for WT Arrays (Affymetrix).

Quality control and differential gene expression was conducted using Transcriptome

Analysis Console (TAC v.4.0.1) software (Affymetrix). Genes were considered differentially expressed if FDRq <0.05. Transcriptomic data subset using genes from Wikipathways “oxidative phosphorylation” or “electron transport chain”, and visualized by hierarchical clustering and principle component analysis using R. Digital cytometry was performed using CIBERSORTx (28), with previously identified mouse specific immune cell signatures (29). Differences in cell fractions were determined by t-test.

### Gene Set Enrichment Analyses (GSEA)

Affymetrix transcriptomic profiling data was exported using TAC, and GSEA (30) was conducted using the Hallmark gene sets (31). FDRq <0.05 was considered significant. For M-Wnt tumors GSEA was also conducted using the Gene Ontology Biological Processes, significant enriched/suppressed gene sets (FDRq <0.05) were then visualized by enrichment mapping (32).

METABRIC mRNA data were accessed via cBioPortal and single sample ssGSEA (ssGSEA) analysis was performed on all patients using the gene set ‘GOBP_HYPOXIA_INDUCIBLE_FACTOR_1ALPHA’. Patients were then grouped into quartiles of ssGSEA scores. The PC expression from Q1 (lowest ssGSEA scores) and Q4 (highest ssGSEA scores) was visualized, and Mann-Whitney U test was performed.

### Normal vs. cancer analysis

Normal and tumor expression levels of PC across multiple tissues were obtained from the Gene Expression database of Normal and Tumor tissues (GENT2) data base (33), and analyzed by Mann-Whitney U test.

### Immunohistochemistry

Rehydrated 5 µm FFPE sections underwent antigen retrieval for 1 hour in a steamer using pH 6.0 citrate buffer, and then progressively blocked with H_2_O_2_ for 10 min and blocking buffer (1% BSA with 5% goat-serum, TBS pH 7.6) for 10 min. Primary antibody incubation occurred overnight at 4°C using anti-CD4 (1:200, Cell Signaling #25229) or anti-CD8 (1:200, Cell Signaling #98941S) antibodies in blocking buffer. Biotinylated secondary anti-rabbit and anti-mouse antibodies (Biolegend Inc. #406401 and #405303) were applied for 1 h and then incubated with ABC (Vectastain, Vector Laboratories #PK-6100) for 30 minutes. Finally, samples were incubated with DAB (Vector Laboratories #SK-4100) for 5 minutes and counterstained with hematoxylin, before being dehydrated and mounted. Visualization of the samples were performed with a light microscope (Nikon Eclipse TS100, Germany) at 20X magnification and positive staining quantification was performed by ImageJ (imagej.nih.gov/ij/download.html, MD, USA). At least 6 randomly selected fields per slide were randomly selected and evaluated by two microscopists one of whom was blinded to groups. The mean positive cell count for each slide was then used for analysis.

### Metabolomics Analyses

E0771 cells were seeded at a density of 2×10^5^ cells/plate in treatment media. The next day, metabolites were extracted using acetonitrile/methanol/water and analyzed by the Metabolomics Center at the University of Illinois Urbana Champaign. Hentriacontanoic acid was added to each sample as an internal standard prior to derivatization.

Metabolite profiles were acquired using an Agilent GC-MS system (Agilent 7890 gas chromatograph, an Agilent 5975 MSD, and an HP 7683B autosampler). The spectra of all chromatogram peaks were evaluated using the AMDIS 2.71 and a custom-built database with 460 unique metabolites. All known artificial peaks were identified and removed prior to data mining. To compare between samples, all data were normalized to the internal standard in each chromatogram, and each sample expressed as relative to control.

### Cell viability and proliferation assays

E0771 cells were seeded at 2×10^3^ cells/well in a 96-well plate (Corning #3610, ME, USA) into low glucose media overnight, then treated with indicated AZD3965 or syrosingopine (SML-1908, Sigma-Aldrich, Burlington, MA) concentrations. After 24 hours of AZD3965 or syrosingopine treatment, cell numbers were quantified by CellTiter-Glo (Promega, WI, USA). Viability was calculated relative to the untreated control. All experiment conditions had six technical repeats and experiments were repeated at least three times.

M-Wnt cells were seeded at 2.5×10^3^ cells/well in a 96-well plate in HPLM overnight, then treated with 25 µM FX-11 (MedChemExpress, HY16214, Monmouth Junction, NJ). After 24 h of FX-11 treatment relative cell numbers were quantified by staining with 100 µl of MTT solution (0.5 mg/mL 3-(4,5-dimethylthiazol-2-yl)-2,5-diphenyl tetrazolium bromide in HPLM without FBS) for 90 min and solubilized in 100 µl DMSO. Cytotoxicity was calculated relative to untreated control.

### Hypoxia

Hypoxic cell culture was achieved by displacing atmospheric O_2_ in a humidified modular incubator chamber (Billups-Rothenberg, Del Mar, CA, #MIC-101) with at least 100 L of 1% O_2_, 5% CO_2_, and 94% N_2_ gas mixture and then incubating at 37°C for 48 hours.

Normoxic control cells were cultured at atmospheric O_2_ at 37°C with 5% CO_2_.

### Western blotting

Western blotting was performed using 20 µg of protein isolated in RIPA buffer, resolved on polyacrylamide gels, and transferred to 0.45 µm PVDF membrane. After blocking for 1 h in 1% BSA/TBST buffer, the membranes were incubated overnight at 4°C with anti-PC antibody (1:500, HPA043922, Millipore-Sigma, Darmstadt, Germany), anti-beta tubulin antibody (1:2000, E7, Developmental Studies Hybridoma Bank (DSHB)), anti-LDH-A (1:1000, 2012S, Cell Signaling Inc., MA, USA), or anti-MCT-1 (1:1000, ab93048 Abcam, Inc., MA, USA).

### Lactate-Glo Assay

The Lactate-Glo assay (Promega, Wisconsin, USA) was used to detect intracellular and extracellular lactate levels *in vitro* and *in vivo*. For the *in vitro* analyses, E0771 and M-Wnt cells were seeded at a concentration of 5×10^3^ cells/well in 96-well plates overnight, then the media was exchanged for treatment media. After a 24 h incubation with MCT-1 inhibitors or FX-11, luminescence values were measured according to the manufacturer’s instructions. Ex vivo tumor lactate was determined by homogenization of 20 mg of tumor in PBS followed by centrifugation (10^4^ RCF, 4°C, 5 min). Lactate levels were determined by Lactate-Glo assay, normalized by total protein, and expressed as relative to scram.

### Extracellular Flux Analysis

Seahorse Metabolic Flux Analyzer XFe96 or XFe24 instruments (Agilent Seahorse Technologies, Santa Clara, CA) were used to determine cellular oxygen consumption rate (OCR) of in vitro scram and PC-suppressed M-Wnt, D2.A1, and E0771 cells. Cells were seeded into Seahorse cell culture plates at a density of 5×10^3^ M-Wnt cells/XFe96 or 3×10^4^ E0771 or 2×10^4^ D2.A1 cells/XFe24 well overnight. 1 hour prior to analysis, cells were incubated at 37°C in assay media (serum-free RPMI-1640 media with 10mM glucose, 2mM glutamine, and 1mM pyruvate, without bicarbonate, pH 7.4) at atmospheric CO_2_. Oligomycin (1.0µM), carbonyl cyanide-4-(trifluoromethoxy)phenylhydrazone (FCCP; 1.0 µM), and rotenone/antimycin A (0.5 µM) were injected sequentially. OCR data was normalized to total protein using a bicinchoninic acid protein assay (Thermo Fisher, Waltham, MA), and then expressed as relative to control.

### High-Resolution Respirometry

An established substrate-uncoupler-inhibitor-titration (SUIT) protocol for high-resolution respirometry (HRR) was utilized with in vitro control and ShPC cells (SUIT-001 O2 ce-pce DOO4, https://www.bioblast.at/index.php/SUIT-001_O2_ce-pce_D004). Cells were re-suspended in Mir05 buffer (Oroboros Instruments, Innsbruck, Austria). 1,000,000 cells were injected into 2 mL pre-calibrated Oxygraph-2k chambers containing Mir05 (O2k, Oroboros Instruments, Innsbruck, Austria). All experiments were performed at 37°C under constant stirring with oxygen concentrations maintained between 100 and 200 µM. Reoxygenation, as needed, was performed via addition of 5 µl of catalase (112,000U/mL dissolved in Mir05, Sigma C9322) and titration of hydrogen peroxide (50 wt. % in H_2_O, Sigma 516813) until desired oxygen concentration was reached. Residual oxygen consumption (ROX) was measured after permeabilization with digitonin and subtracted from oxygen flux as a baseline for all respiratory states to obtain mitochondrial respiration. Specific flux was expressed as oxygen consumption per million cells (*ρ*mol.s^-1^.million cells). To determine flux control ratios (FCR), respiration values for each replicate were normalized to its respective NS_E_ specific flux. DatLab Software (V7.4, Oroboros Instruments) was utilized for data acquisition and post-experimental analysis

### Statistical Analyses

Python 3.8.5, R 4.0.2, Graphpad Prism 8 software was used for statistical analysis (GraphPad Software Inc., La Jolla, CA, USA, www.graphpad.com). R packages used were FactoMiner 2.4, factoextra 1.0.7, gplots 3.1.1, and ggpubr 0.4.0. One- and two-way ANOVAs with Tukey’s post-hoc testing, were used where three or more groups existed and t-tests were used to compare two groups. Welch’s correction was applied where variance was not similar between groups. P-values of less than 0.05 were considered significant. Where multiple comparisons were made p values were considered significant where FDRq < 0.05. No exclusion criteria were utilized in these studies.

### Data availability

All transcriptomic data has been deposited to GEO, M-Wnt tumor transcriptomic data is available at GSE199828, E0771 tumor transcriptomic data is available at GSE199829, and E0771 in vitro transcriptomic data is available at GSE199830. All other data is available on reasonable request to the corresponding authors (SDH and MKW).

## RESULTS

### A dichotomous role for PC in lung metastasis versus primary tumors

Our previous studies utilized several BALB/c syngeneic models of breast cancer to demonstrate that PC is required for metastatic colonization of the lungs (24). To extend these observations we similarly conducted tail vein injection assays in C57BL/6J mice using syngeneic E0771 mammary tumor cells with or without knockdown of PC expression (**Fig. 1A-C**). Consistent with our previous work, the development of lung metastasis following tail vein injection was attenuated in PC-suppressed E0771 cells relative to scramble control (**Fig. 1D-H**). To investigate the impact of PC suppression on primary tumor growth, scrambled control and PC-suppressed cells were orthotopically transplanted into the 4^th^ mammary fat pad (**Fig. 1I**). In contrast to the inhibitory effect of PC suppression on metastatic seeding of the lungs following tail vein injection, orthotopic tumors from PC-suppressed E0771 cells were larger than the scrambled control (**Fig. 1J**). No difference in spontaneous metastatic burden was found between PC-suppressed or scrambled control tumors (**Fig. 1K**).

**Figure 1:**
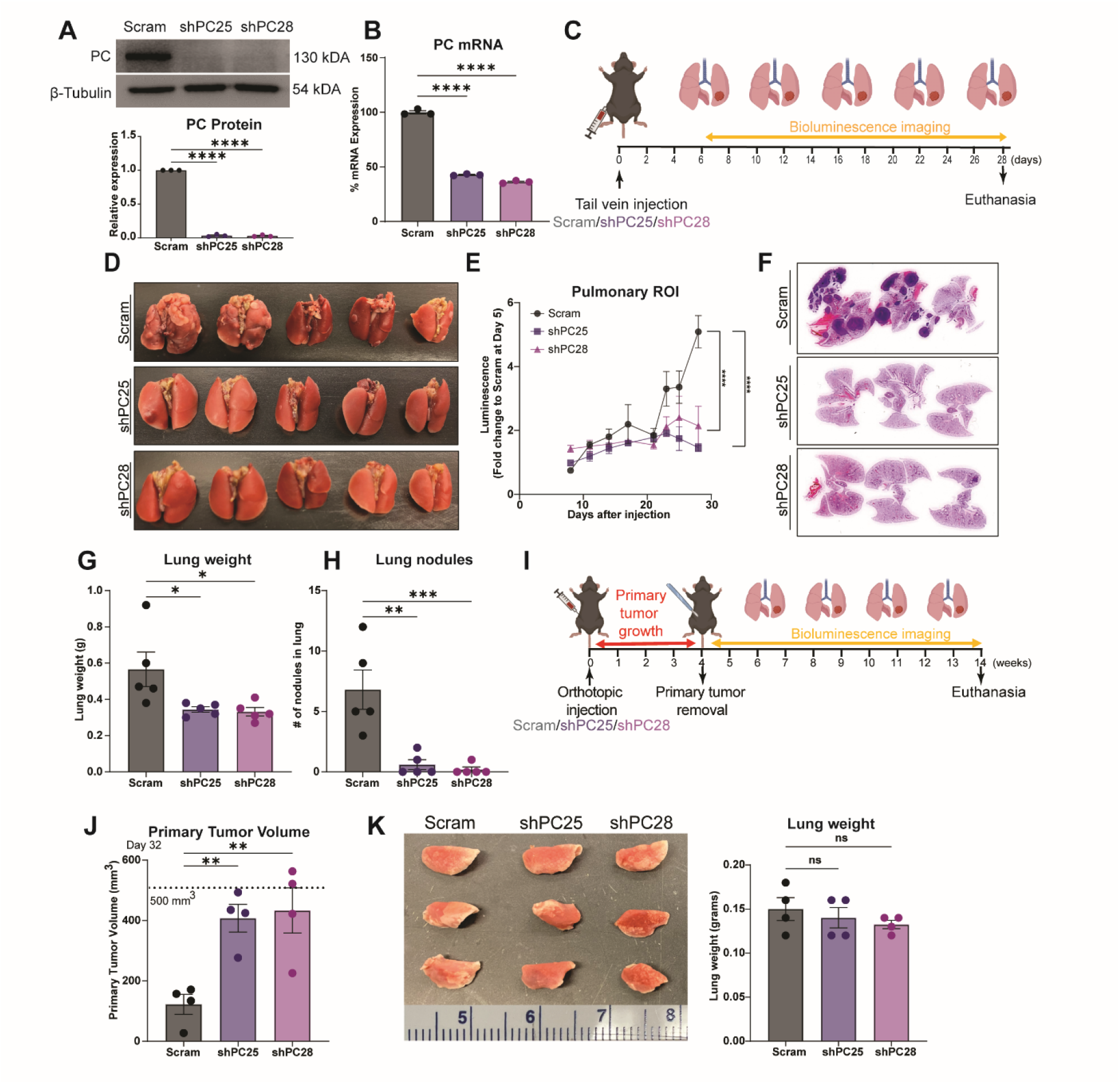
A dichotomous role for PC in metastatic pulmonary and primary tumors. (A) Immunoblot analyses of control (scram) and PC-suppressed (shPC25 and shPC28) E0771 cells probed for PC and ß-tubulin. (B) Expression of PC in control and PC-suppressed E0771 cells was quantified by qPCR (n=3/group). (C) Experimental timeline of E0771 injected C57BL6/J mice. Mice were injected via the lateral tail vein with 10^6^ E0771 cells. Progression of pulmonary tumors was tracked by bioluminescence imaging every other day for 28 days. (D) Lungs from mice injected with control and PC-suppressed E0771 cells. (E) Pulmonary luminescence of the 3 groups (5/group) of mice between Day 8 and 28 following tail vein inoculation. (F) H&E staining of lung histological sections (n=3/group) (G) Weight and (H) pulmonary tumor nodule counts at euthanasia (5/group). (I) Experimental timeline where 5×10^5^ control and PC-suppressed E0771 cells orthotopically transplanted into C57BL6/J mice (5/group). After 4 weeks, primary tumors were removed. Progression of pulmonary metastasis was tracked by bioluminescence imaging for another 10 weeks and animals were sacrificed. (J) Primary tumor volume measurements of control and PC-suppressed tumors (n=4/group) (K) Images and weights of lungs at euthanasia (n=4/group). Statistical significance determined by one-way ANOVA (*p<0.05; **p<0.01; ***p<0.001; ****p<0.0001).

To further investigate the role of PC on orthotopic mammary tumor growth, we utilized M-Wnt cells, another C57BL/6 syngeneic tumor model. Using these cells, we established doxycycline-inducible and constitutive models of PC suppression. Orthotopic tumors from constitutive PC-suppressed M-Wnt cells were larger than scrambled control tumors (**Fig. 2A**). Consistent with constitutive PC-suppression in E0771 and M-Wnt cells, suppression of PC by doxycycline treatment resulted in significantly larger tumors relative to non-doxycycline treated control mice following orthotopic injection of M-Wnt cells with a doxycycline-inducible shPC construct (**Fig. 2B-C**). To identify major pathways and processes disrupted by suppression of PC, we conducted global transcriptomics using an Affymetrix microarray followed by gene set enrichment analysis (GSEA). We found striking suppression of GSEA Hallmark gene sets related to immune signaling in the shPC group relative to control M-Wnt tumors (**Fig. 2D**). To identify more granular pathway alterations, we next conducted GSEA using Gene Ontology Biological Processes gene sets with enrichment mapping to minimize redundancy. We found that relative to control tumors, PC-suppressed tumors had disrupted immune-related and fatty acid/lipid metabolism signaling (**Fig. 2E**).

**Figure 2:**
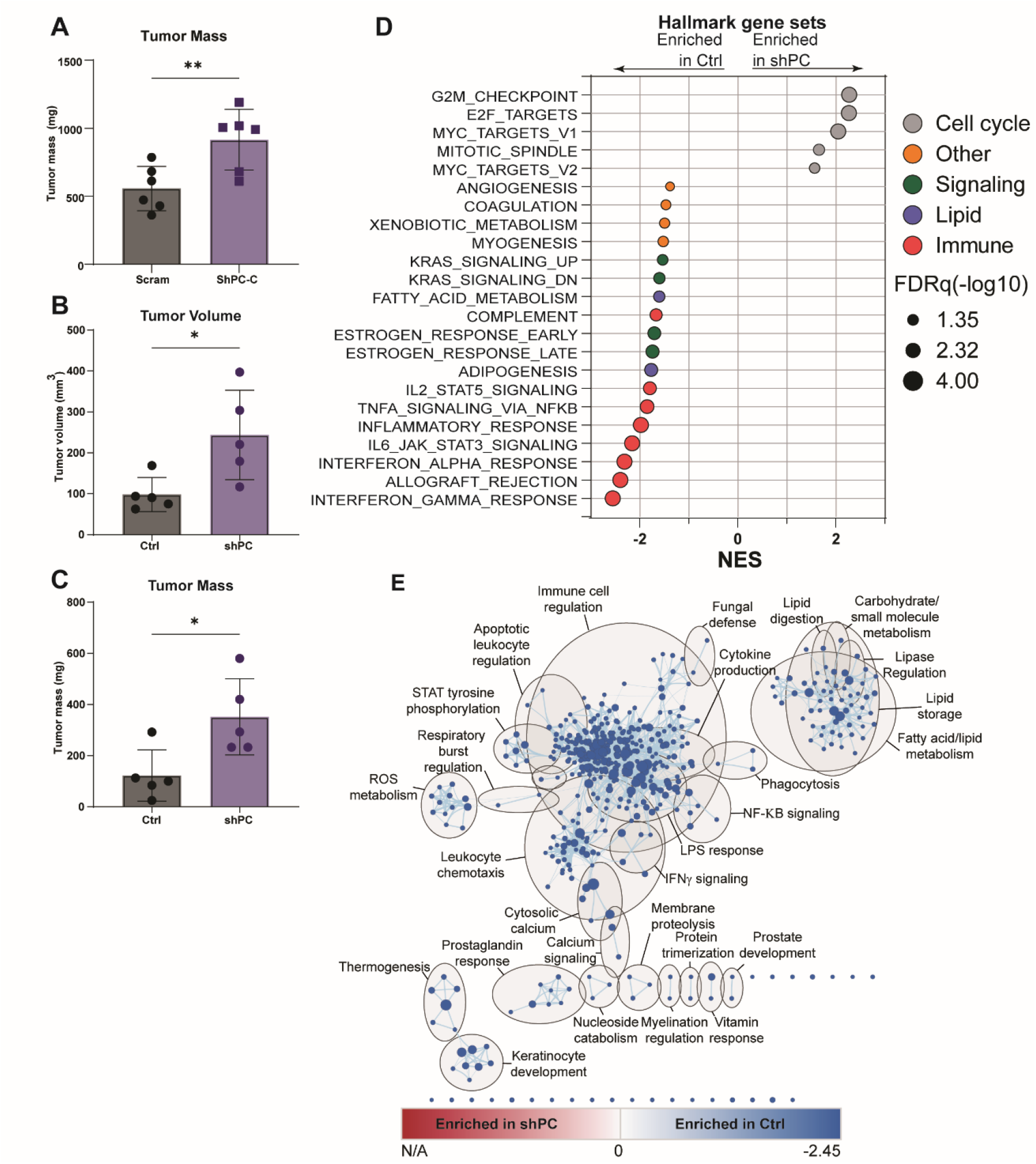
Suppression of PC increases primary tumor growth and suppresses immune related gene expression. ((A) Tumor mass from M-Wnt tumors harboring control (scram) or PC-targeted (ShPC-C) constitutive shRNAs (n=6/group). (B) Tumor volume and (C) tumor mass from M-Wnt tumors harboring PC-targeted dox inducible shRNA treated with (ShPC) or without doxycycline (Ctrl) (n=5/group). (D) Hallmark GSEA of transcriptomic profile of tumors from B-C, which were significantly enriched with FRDq <0.05 (n=5/group). (E) Enrichment maps of significant (FDRq <0.05) GSEA Gene Ontology Biological processes in Ctrl and shPC tumors. (FDRq<0.05). Node color indicates normalized enrichment score, node size indicates gene set size, line weight indicates degree of overlap, and clusters indicate minimum 50% overlap of gene sets. Statistical significance was determined by Student’s t-test (A-C) (*<0.05, *<0.01).

We next conducted digital cytometry to identify potential immune cell populations that were altered by suppression of PC and identified an increase in M0 macrophages and a reduced abundance of Th1 and resting NK cells (**Fig. S1A-E**). Finally, we confirmed these effects in orthotopic E0771 tumors by performing transcriptomic analysis followed by GSEA using Hallmark gene sets on scrambled control and PC-suppressed E0771 tumors. We once again found that PC suppression in tumor cells suppressed tumor immune-related signaling relative to control tumors (**Fig. S2**). Overall, these data indicate that unlike the essential role of PC in lung metastasis (24), suppression of PC may aid in primary tumor development by modulating the tumor immune microenvironment.

### PC expression is reduced by hypoxia

To investigate the prevalence of PC suppression in primary tumors we used data publicly available through GENT2 to compare PC mRNA levels in normal tissues to corresponding primary tumors from patients from either any cancer, or specifically breast or lung cancer. PC expression was significantly lower in primary tumors relative to normal tissue in both the pan-cancer and breast cancer data sets (**Fig. 3A**). In contrast PC expression was higher in lung cancer relative to normal (**Fig. 3A**). Given the key role of PC in directing central carbon metabolism, and the prevalence of hypoxia in solid tumors, we next sought to determine if PC expression was correlated with markers of hypoxia. Gene expression data from patients with breast cancer was obtained from METABRIC and stratified into quartiles based on ssGSEA enrichment of HIF-1α signaling and PC expression was determined from RNAseq data.

**Figure 3:**
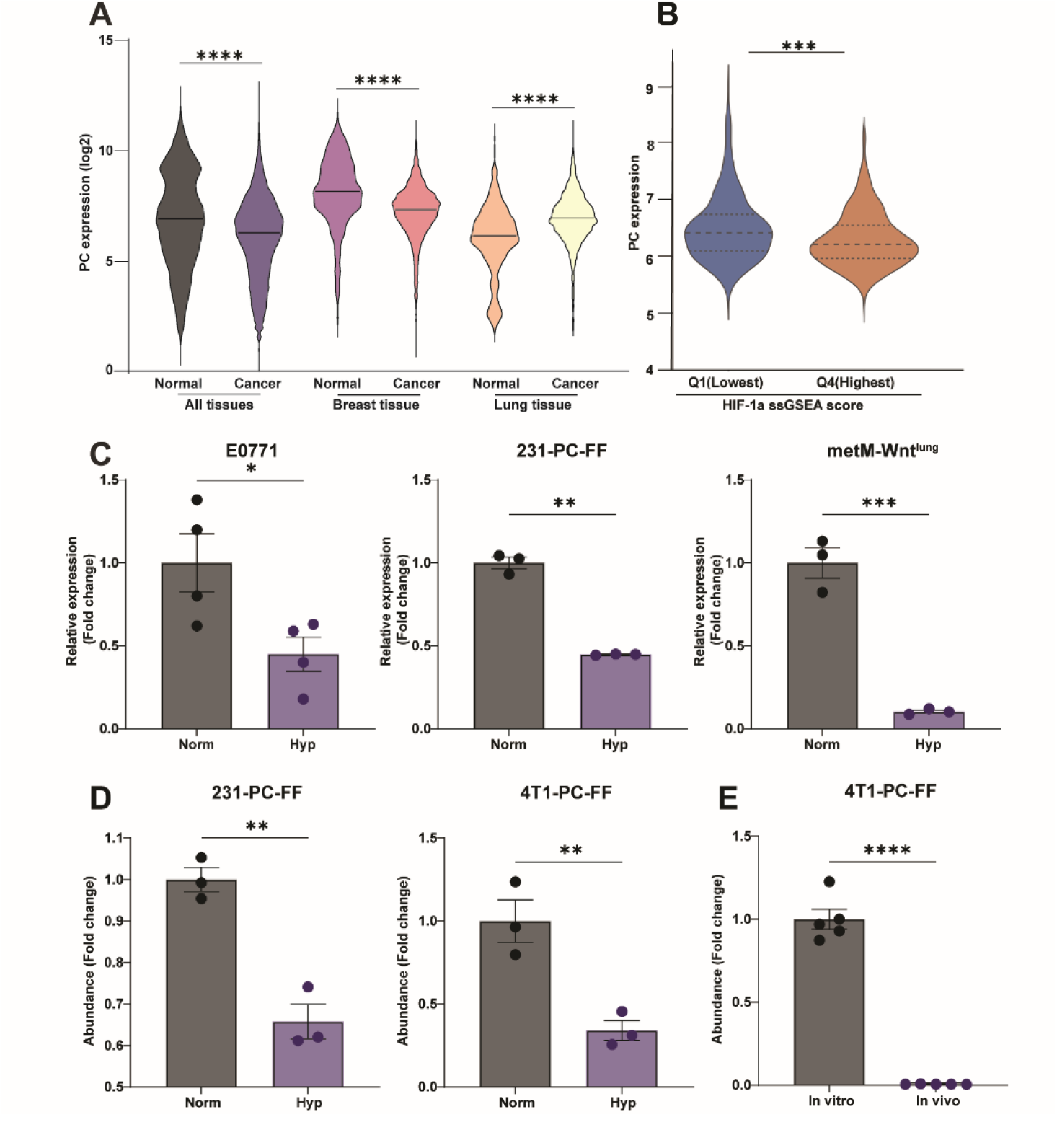
PC expression is reduced by hypoxia. (A) Comparison of PC expression profiles in publicly available databases among all tissues (n=5487 normal, 35806 cancer), breast (n=475 normal, 5555 cancer), and lung tissues (n=1016 normal, 2316 cancer). (B) PC expression in and upper and lower quartiles of HIF-1α signaling ssGSEA score in the METABRIC database. (C) PC-gene expression determined by qPCR under normoxia (Norm) and hypoxia (Hyp) in E0771, MDA-MB-231 and metM-Wnt^lung^ cells (n=3-4/group). (D) MDA-MB-231 (231) cells transiently transfected with and 4T1 cells stably expressing firefly luciferase under the control of the PC promoter and renilla luciferase under the control of the CMV promoter cultured under normoxic (Norm) and hypoxic (Hyp) conditions and analyzed using the Dual-Glo Luciferase Assay System (n=3/group). (E) PC promoter luciferase reporter 4T1 described in panel E orthotopically transplanted for 2 weeks and *ex vivo* PC reporter activity compared to *in vitro* cultured cells (n=5/group). Statistical significance was determined by Mann-Whittney U test (A-B) or Student’s t-test (C-E) (*p<0.05;**p<0.01, ****p<0.0001).

We found that PC expression was significantly lower in tumors that were enriched for this hypoxia signature (**Fig. 3B**). Hence, we sought to determine if hypoxia could directly suppress PC expression. Using several models of breast cancer in vitro, we found that cells cultured under hypoxic conditions reduced levels of PC mRNA relative to normoxic conditions (**Fig 3C**). All three cell types demonstrated 50-90% reduction in PC mRNA levels after 48-hours of culture under hypoxic conditions (**Fig. 3C**). Previous studies identified a HIF-1α binding site in the PC proximal promoter (34). Hence, we cloned the PC proximal promotor, from -800 to +61 relative to transcriptional start, upstream of firefly luciferase to determine promotor activity under normoxic and hypoxic conditions. Hypoxia reduced reporter activity in MDA-MB-231 and 4T1 cells relative to normoxia (**Fig. 3D**). Orthotopic 4T1 tumors are extremely hypoxic (35), and our previous studies indicate PC is not detectable in 4T1 primary tumor cells (24). Hence, we tested if growth in such a hypoxic environment might reduce PC expression by reducing PC promotor activity. Orthotopic tumors generated with 4T1 cells stably expressing the PC reporter had lower PC promoter activity relative to in vitro cultured cells (**Fig. 3E**). Taken together, these data indicate that PC expression is reduced by hypoxic conditions that are characteristic of large solid tumors.

### Suppression of PC increases lactate production

We next sought to elucidate potential mechanisms through which PC suppression may promote an immunosuppressed microenvironment. We first conducted metabolomic analyses on control and PC-suppressed E0771 cells, which revealed that PC regulates levels of both glycolytic and TCA cycle intermediates. As expected, levels of oxaloacetate, the immediate product of PC-mediated carboxylation of pyruvate, were decreased in PC-suppressed cells relative to control (**Fig. 4A**). However, most other TCA cycle intermediates had increased pool sizes in PC-suppressed cells relative to control (**Fig. 4A**). Further, we observed a 10-fold increase in intracellular lactate in PC-suppressed E0771 cells as compared with control (**Fig. 4A**). Concordant with dysregulated TCA cycle metabolism, transcriptomic profiling revealed marked up-regulation of several transcripts encoding TCA enzymes in PC-suppressed E0771 cells (**Fig. 4B**). We next assayed extracellular lactate levels and also found elevated levels in PC-suppressed E0771 and M-Wnt cells relative to controls (**Fig. 4C-D**). Consistent with increased lactate production and secretion, immunoblot analyses indicated that PC-suppressed cells express higher levels of LDH-A and MCT-1 compared with control cells (**Fig. 4E**). In addition, PC-suppressed E0771 cells were more sensitive to the MCT1 inhibitor AZD3965 (**Fig. 4F**), with similar effects seen with the MCT1/MCT4 dual inhibitor syrosingopine (**Fig. 4G**). Abrogation of the PC-mediated lactate secretion demonstrated the effectiveness of 200nM AZD3965 to inhibit MCT-1 (**Fig. 4H**). Finally, we confirmed that suppression of PC expression increased the sensitivity of M-Wnt cells to the LDHA inhibitor FX-11, and that increased lactate production was reverted by FX-11 treatment (**Fig. S3A-C**).

**Figure 4:**
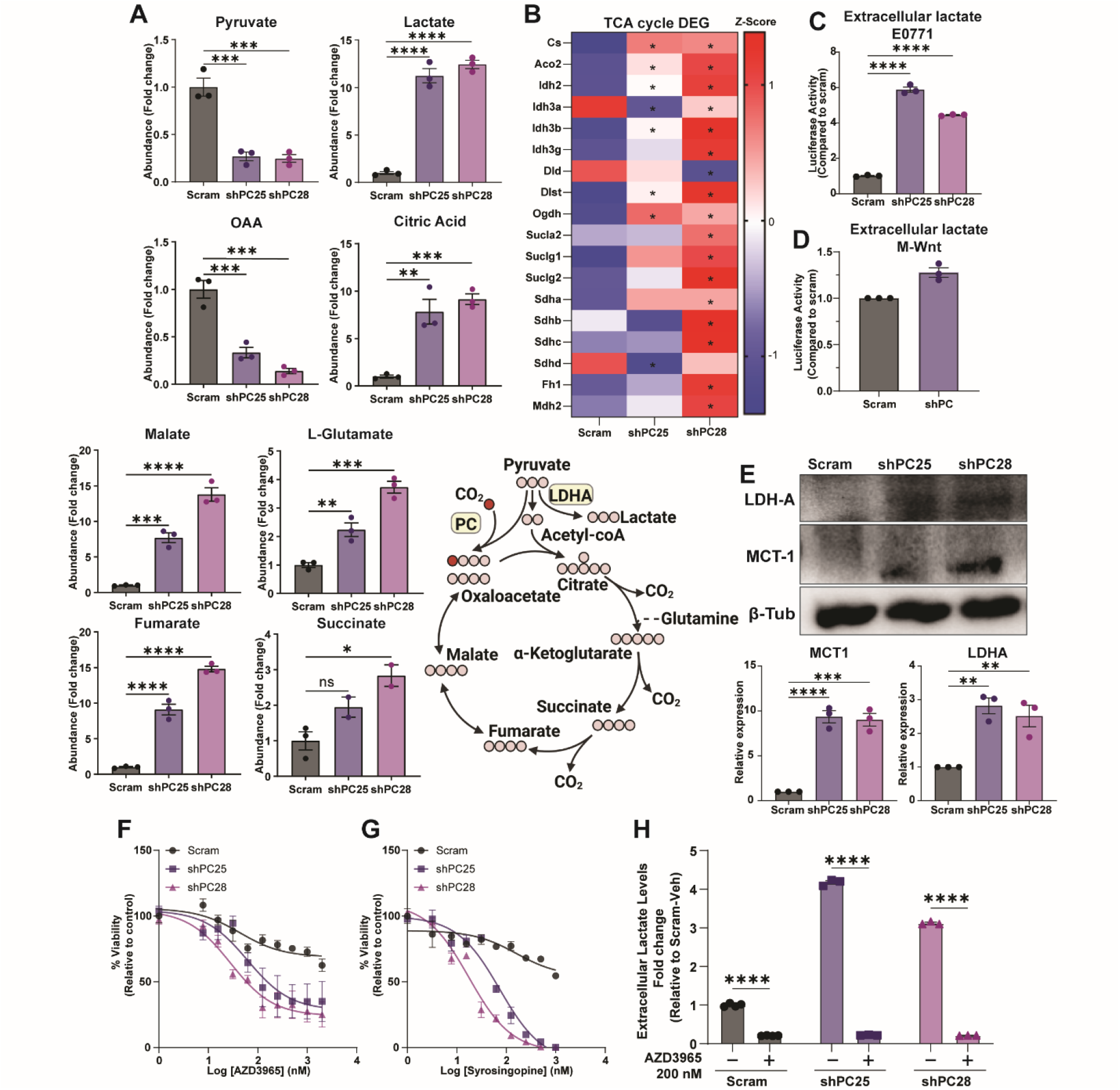
Suppression of PC increases lactate production. ((A) Comparison of glycolytic and TCA cycle intermediates between control and PC-suppressed E0771 cells (n=3/group). (B) Heatmap of differentially expressed TCA cycle genes determined by transcriptomic profiling (FDRq <0.05) (n=3/group). (C-D) Extracellular lactate concentration in control and PC-suppressed E0771 and M-Wnt cells as quantified by luminescence assay. (E) Immunoblot analyses for LDH-A and MCT-1 in control and PC-suppressed cells. (F-G) Cell viability analyses of control and PC-suppressed E0771 cells upon treatment with the MCT-1 inhibitors, AZD3965 (n=3/group), or the MCT 1 and 4 inhibitor syrosingopine (n=2/group). (H) Extracellular lactate levels quantified following treatment of control and PC-suppressed E0771 cells with 200nM AZD3965 (n=3/group). Statistical significance was determined by one-way ANOVA (A, C, D) or two-way ANOVA (F-H) (*p<0.05; **p<0.01; ***p<0.001; ****p<0.0001). phosphorylation” gene list curated by Wikipathways, demonstrated distinct clustering of cell lines (**Fig 5A-D)**. To characterize metabolic differences between control and PC-suppressed cells, we next performed extracellular flux analysis. PC suppression in three

Using transcriptomic profiling of PC-suppressed E0771 cells in vitro, we next sought to identify metabolically related pathway alterations. PCA and hierarchical clustering of gene expression, using the member genes of “electron transport chain” or “oxidative different mammary cancer cell lines decreased oxygen consumption rate (OCR) compared with their PC-expressing counterparts (**Fig 5E-G**).

**Figure 5:**
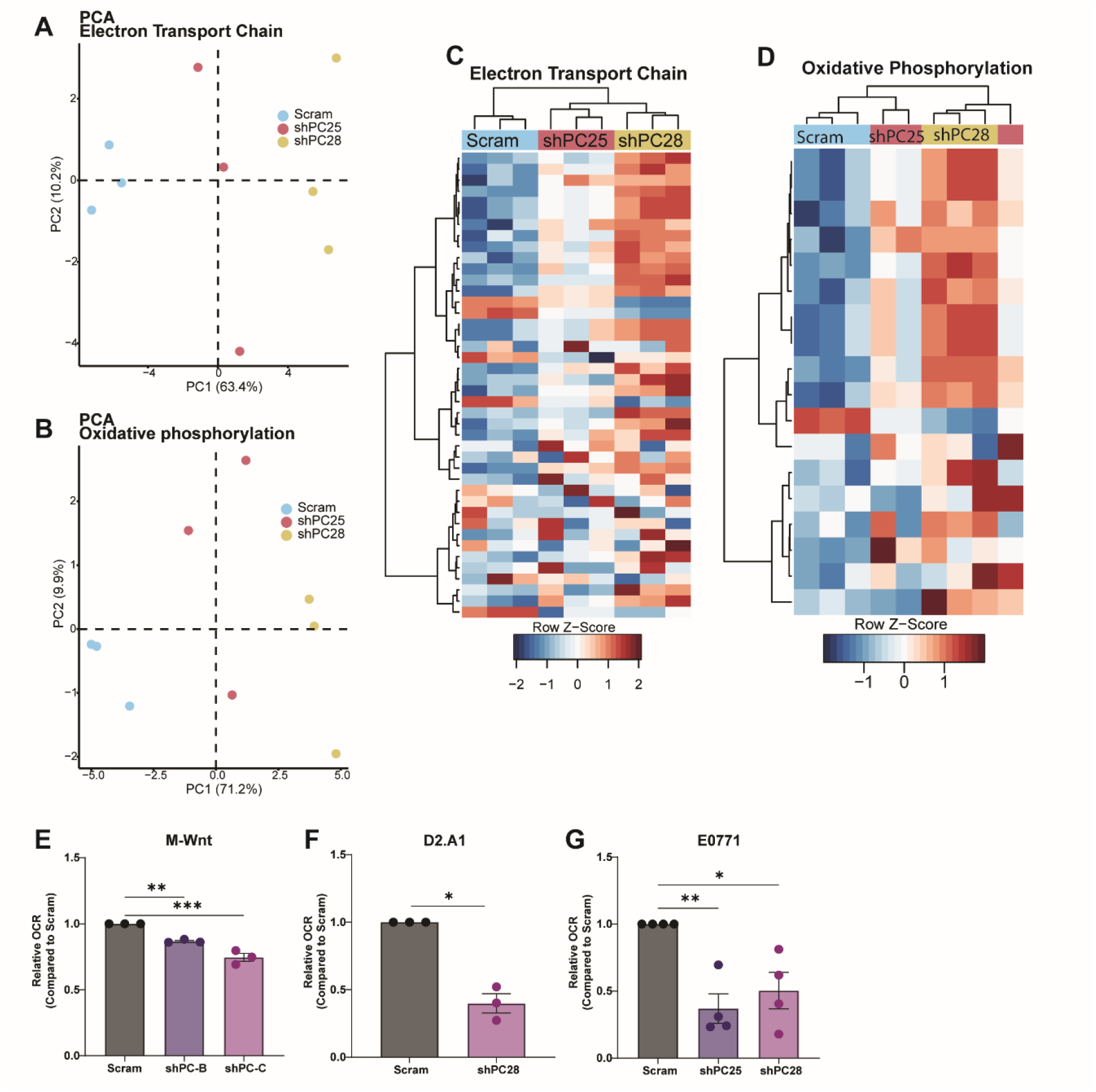
Suppression of PC decreases oxygen consumption. (A-B) PCA of gene expression profiles of control (scram) and PC-suppressed (ShPC25 and ShPC28) E0771 cells cultured in low (5.6 mM) glucose DMEM using genes contained in the “Electron transport chain” (A) and “Oxidative phosphorylation” Wikipathways (n=3/group). (C-D) Hierarchical clustering of gene expression profiles obtain under conditions described in panels A and B. (E-G) Basal oxygen consumption rates determined by extracellular flux analysis of the indicated control and PC-suppressed cells(n=3-4/group). Statistical significance calculated using Student’s t-test (*<0.05, **<0.01).

To determine if the reduction in OCR observed in PC-suppressed cells was due to electron transport chain intrinsic defects, we used high resolution respirometry to measure mitochondrial OCR in the presence of non-limiting ETC substrates. Reduced O_2_ consumption was not found with any combination of complex I and II substrates tested following suppression of PC in M-Wnt cells. Indeed, shPC-B showed an increased in complex I and II mediated O_2_ consumption (**Fig. S4A**). Similarly, when O_2_ consumption for each sample at each stage in the SUIT protocol were normalized to peak O_2_ consumption (PMGS_E_), no alterations in flux control ratio were observed (**Fig. S4B**). Together, these data suggest that suppression of PC impairs TCA metabolism and oxidative phosphorylation independent of electron transport chain intrinsic defects.

### Tumor microenvironment immunosuppression following suppression of PC requires lactate secretion

We next sought to determine if suppressing PC would alter the abundance of tumoral CD4^+^ and CD8^+^ T cells. We quantified CD4^+^ and CD8^+^ lymphocytes in tumors from control and PC-suppressed groups by immunohistochemistry. Consistent with our gene expression analyses, we observed lower numbers of CD8^+^ lymphocytes and higher numbers of CD4^+^ T cells in PC-suppressed E0771 tumors (**Fig. 6A**). To determine if extracellular lactate mediates intratumoral lymphocyte abundance, we treated both control and PC-suppressed tumor bearing mice with the MCT-1 inhibitor, AZD3965, every day for two weeks (**Fig. 6B**). MCT-1 inhibitor treatment significantly reduced tumor growth in both control and PC-depleted groups, but significant accumulation of intracellular lactate was only observed in the PC-depleted tumors (**Fig. 6C-D**).

**Figure 6:**
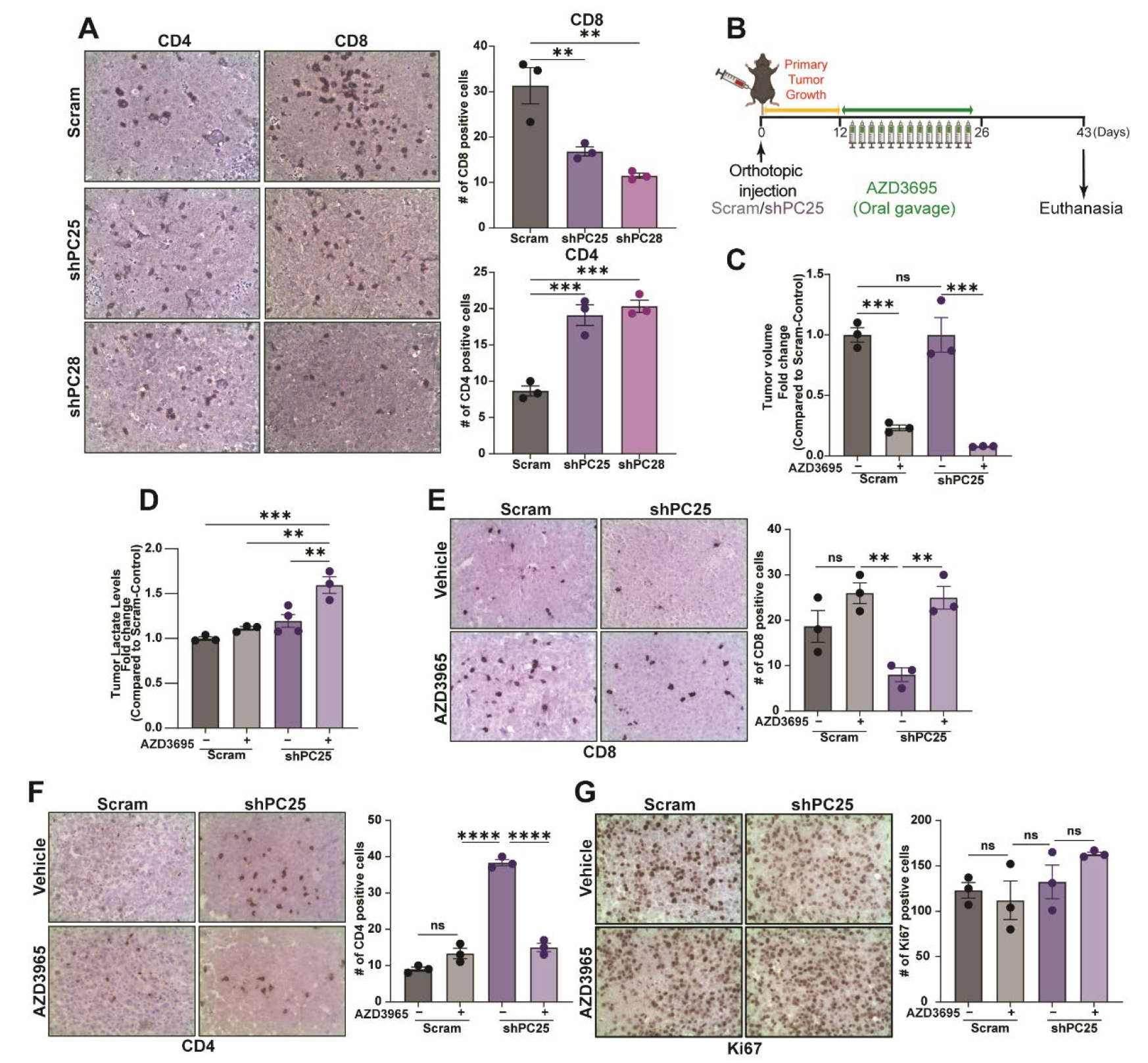
Suppression of PC modulates the tumor immune microenvironment. (A) (left panel) Representative IHC staining of sections from control and PC-suppressed E0771 tumors. (Right panel) Quantification of CD8^+^ and CD4^+^ T cell numbers (n=3/group). (B) Experimental summary of the *in vivo* study. Mice were orthotopically engrafted with control and PC-suppressed E0771 cells (5×10^5^ cells per mouse) and their primary mammary tumor growth was observed for 10 days at which point AZD3965 was administered (C) Fold change of tumor volume in response to AZD3965 (n=3/group). (D) Lactate levels in the indicated tumors determined by luminescent assay (n=3-4/group). (E-G) (left panels) Representative IHC for CD8, CD4, and Ki67 staining of tumor sections from control and PC-suppressed E0771 tumors under control and AZD3965 treated conditions. (Right panel) Quantification of staining CD8^+^ and CD4^+^ T cells, and Ki67^+^ cells (n=3/group). Statistical significance determined by one-way ANOVA (ns: non-significant, **<0.01, ***<0.001, ****<0.0001).

Importantly, immunohistochemical analyses of those tumor samples clearly demonstrated that MCT-1 inhibition mitigate d the effects of tumor cell PC suppression on tumoral CD4^+^ and CD8^+^ lymphocyte abundance (**Fig. 6E-F**). Finally, MCT-1 inhibition did not alter the number of Ki67 positive cells in the tumors of any group, indicating that the antitumor effect of MCT-1 inhibition was not explained by global suppression of proliferation (**Fig. 6G**). These results demonstrate that the modulation of the tumor immune microenvironment by PC-suppression is dependent on lactate secretion.

## DISCUSSION

Cancer cell metastasis and immunosuppression both rely on remodeled tumor cell metabolism (3,4). We and others have shown in several metastatic mammary tumor models that PC is essential in the initiation of pulmonary metastases (23-25). Here we describe a model whereby suppression of PC by hypoxia drives immunosuppression in the TME via extracellular lactate, thereby supporting tumor growth (**Fig. 7**). Concordant with our findings, recent work has identified suppression of PC in TAMs to limit antitumor immunity, with both hypoxia and soluble factors from the TME required for PC suppression (26). Thus, our study builds a framework for understanding the role of PC in primary tumors as a key hypoxia-responsive factor that drives metabolic reprogramming of cancer cells and immunosuppression in tumors.

**Figure 7:**
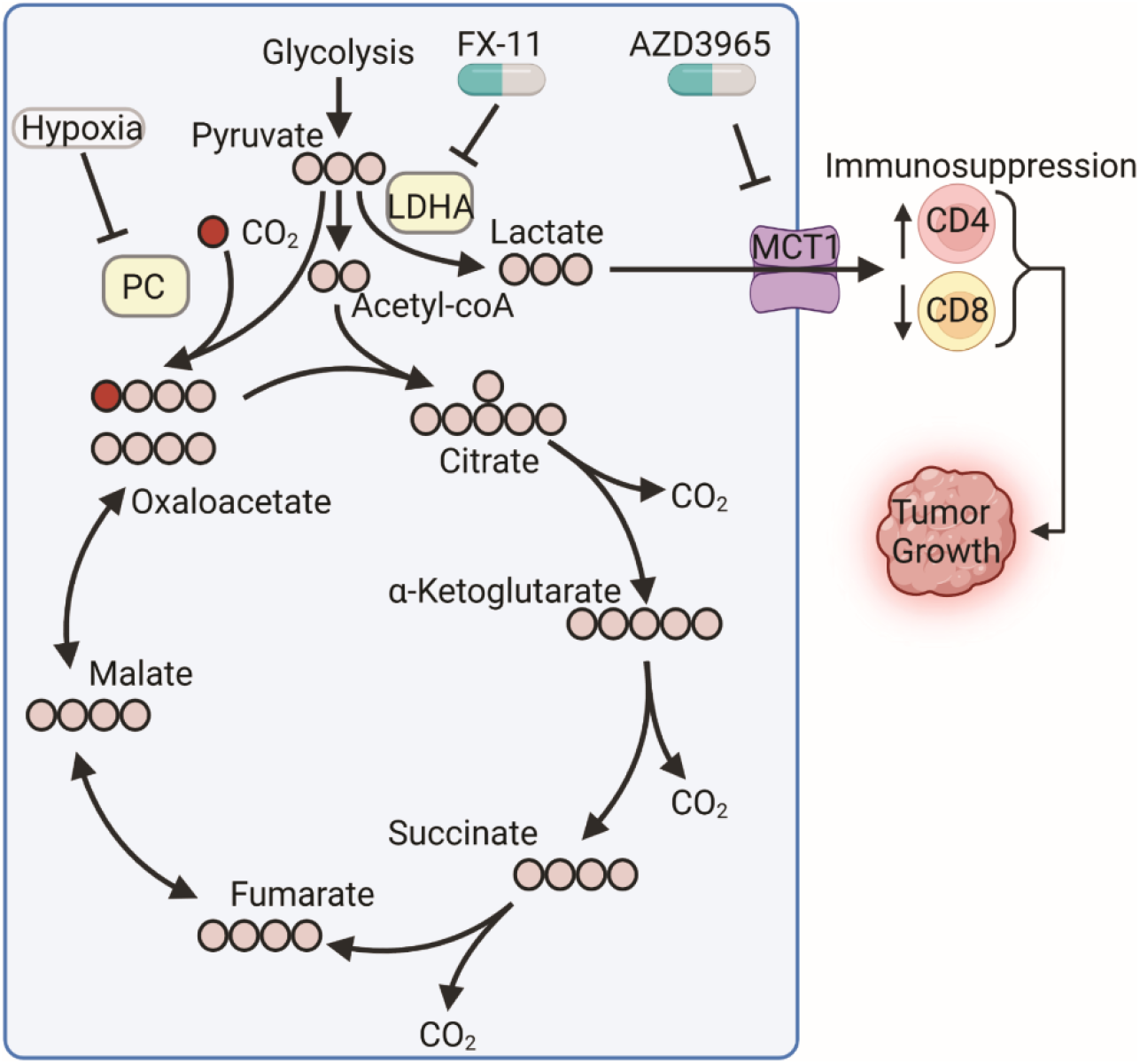
Framework for PC suppression directed immunosuppression. PC expression maintains oxaloacetate pools through carboxylation of pyruvate. Hypoxia potently suppresses PC in cancer cells. Suppression of PC limits oxaloacetate pools and promotes increased lactate production. Increased lactate metabolism promotes immunosuppression in the TME. Lactate metabolism is a targetable dependency in cancer cells following PC suppression.

Hypoxia-mediated changes in tumor metabolism remain incompletely understood (36). HIF-1α is known to limit oxidative phosphorylation via activation of pyruvate dehydrogenase kinase (PDK) (37), and promote lactate metabolism by direct activation of lactate dehydrogenase (LDH) enzymes (38,39) and MCT transporters (40). Our work extends upon these findings by demonstrating that under hypoxic conditions, PC is transcriptionally suppressed via its proximal promoter, and loss of PC promotes increased lactate production. These findings are consistent with patient data that indicate PC expression is lower in many solid tumors relative to normal tissue. Further, PC expression is inversely associated with a hypoxic signature of primary breast tumors. Moreover, our previous studies also indicate that the extremely hypoxic primary tumors resulting from engraftment of 4T1 cells also lack PC expression (24).

To recapitulate the loss of PC expression in aggressive tumors, we utilized various shRNA constructs to deplete PC in E0771 and M-Wnt cells, as they spontaneously metastasize at lower rates than 4T1 cells. Indeed, suppression of PC in both E0771 and M-Wnt cells led to greater primary tumor growth. Metabolomics analysis revealed that PC suppression leads to the expected depletion of oxaloacetate and increased levels of citric acid. Interestingly, under nutrient replete culture conditions we also observed increased levels of glutamate, succinate, fumarate, and malate. These data are consistent with previous reports that indicate higher levels of pyruvate and lower levels of glutamine drive the requirement of PC in the pulmonary microenvironment (25). In contrast, glutamine levels dominate in primary mammary tumors, potentially making PC expression dispensable. Lactic acidosis is a clinical consequence of genetic PC deficiency in newborns (41). These data provide strong clinical precedent for PC insufficiency to increased lactate production. Here, we demonstrate that PC-suppressed cells produce more lactate and are more sensitive to inhibition of lactate metabolism using inhibitors of lactate synthesis and secretion. Taken together these data suggest that PC suppression in primary tumors may drive increased glutamine utilization for anaplerotic metabolism to support rapid lactate efflux.

Suppression of PC also reduced markers of tumor immunosurveillance in both tumor models used in this study. IHC staining further supported these gene expression differences by demonstrating reductions of CD8^+^ T cells and increases of CD4^+^ T cells in PC-suppressed tumors relative to controls. The finding that loss of PC in cancer cells regulates CD8^+^ and CD4^+^ T cell populations in tumors maybe an important advance in understanding the lactate-mediated suppression of antitumor immunity (6,10,16). Our finding that pharmacologically inhibiting (via AZD3965) lactate transport reversed remodeling of tumoral T cell populations is striking as it indicates that not only was loss of PC expression sufficient to promote these effects, but also highlights lactate metabolism as an exploitable dependency. Thus, our current study establishes that PC-mediated changes in the TME are dependent on enhanced lactate secretion by tumor cells.

Taken together, our findings link tumor hypoxia to aberrant production of lactate via suppression of PC. In contrast to the essential role of PC for metastatic spread to the oxygen rich microenvironment of the lungs, the current study presents an important relationship between metabolic plasticity and regulation of tumor immunity. This work fits within a growing body of evidence that supports PC as an important regulator of both antitumor immunity and cancer metabolism.

## Acknowledgments

This work was supported by a grant from the National Institutes of Health (NIH)(R01CA232589) to MKW, DT, and SDH. This research was also supported in part by grants from; the American Cancer Society (RSG-CSM130259) and the NIH (R01CA207751; R21AA026675) to MKW; and the Breast Cancer Research Foundation (BCRF-21-073) and the NIH (R35 CA197627) to SDH. We also acknowledge the support of the Purdue Center for Cancer Research via its NIH grant (P30CA023168).

We kindly acknowledge the expertise of the personnel within the Purdue Center for Cancer Research Biological Evaluation Core. We also acknowledge the use of the facilities within the Bindley Bioscience Center, a core facility of the NIH-funded Indiana Clinical and Translational Sciences Institute.

