## Supplemental figures for "Hypoxia-mediated suppression of pyruvate carboxylase drives tumor microenvironment immunosuppression": supp. figure legends.docx

**Figure S1: Digital cytometry of shPC and Ctrl tumors**

Digital cytometry based on transcriptomic profiler from control and PC-suppressed M-Wnt tumors (n=5/group). (A) M0 Macrophages, (B) M1 Macrophages, (C) naïve CD8 T cells, (D) TH1 cells, and (E) resting NK cells (n=5/group). Statistical significance was determined by Student’s t-test (A-E) (ns: non-significant, * <0.05). All graphs represent mean ± SEM.

**Figure S2: Suppression of PC in E0771 tumors suppresses immune related gene expression**

Hallmark GSEA of transcriptomic profile of tumors from control and PC-suppressed E0771 tumors, significantly enriched with FRDq <0.05 (n=3/group).

**Figure S3: Suppression of PC increases lactate production and sensitizes M-Wnt cells to lactate metabolism inhibition**

(A) Expression of PC in control (scram) and PC-suppressed (ShPC-B) M-Wnt cells was quantified by qPCR (n=3/group). (B) Cell viability analysis of control and ShPC-B cells upon treatment of 25$\mu$M FX-11(n=4/group). (C) Intracellular lactate concentration in control and ShPC-B cells as quantified by using luminescent assay(n=3/group). Statistical significance determined by Student’s t-test (A-B) and one-way ANOVA (C) (**<0.01, ****<0.0001).

**Figure S4:** **PC suppression does not promote electron transport chain intrinsic defects**

High resolution respirometry of permeabilized M-Wnt cells. (A) Specific oxygen flux of control (scram) and PC-suppressed (ShPC-B and ShPC-C) M-Wnt cells ($\rho$mol$\cdot$s^-1^$\cdot$million cells) (n=8/group). (B) Flux control ratio (FCR) normalized to the internal reference state PMGS_E_, achieved following addition of N- and S-coupled substrates and titration of FCCP until complete uncoupling of the mitochondrial membrane potential was achieved. Statistical Significance determined by one-way ANOVA. *p-values < 0.05. **p-values < 0.01
