## Supplementary figures and images for "Hypoxia-mediated suppression of pyruvate carboxylase drives tumor microenvironment immunosuppression"

### SuppFig1.pdf

#### Fig. S1

##### M-Wnt cells

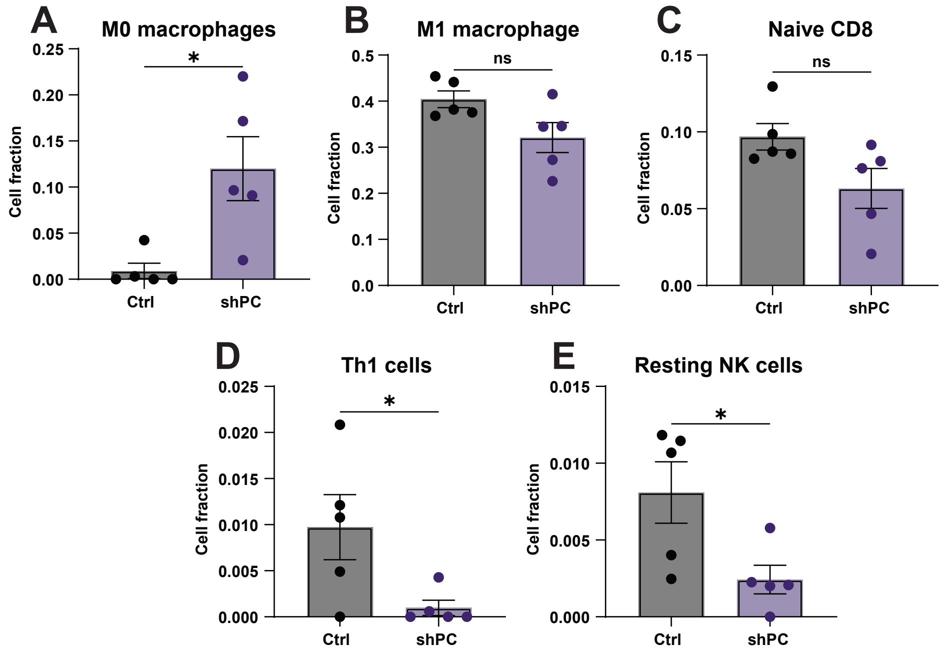

### SuppFig2.pdf

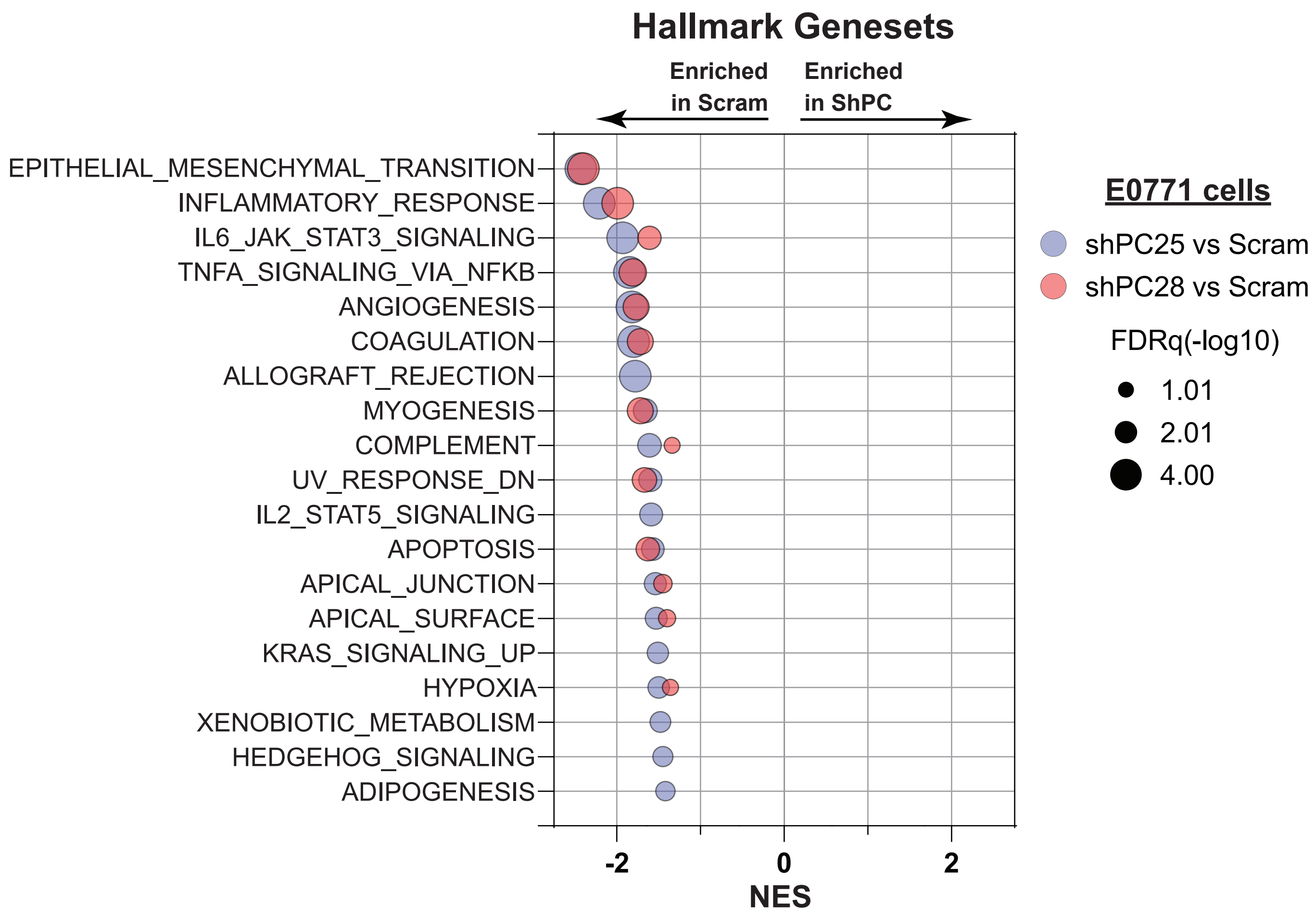

### SuppFig3.pdf

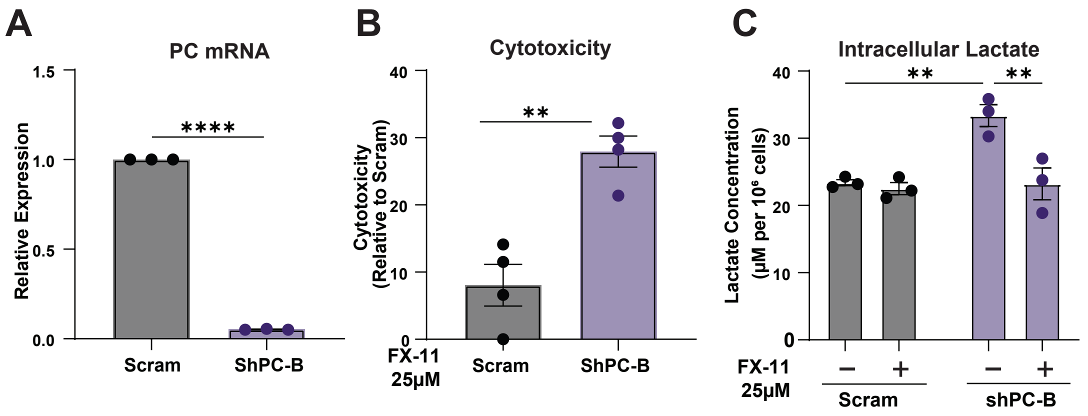

### SuppFig4.pdf

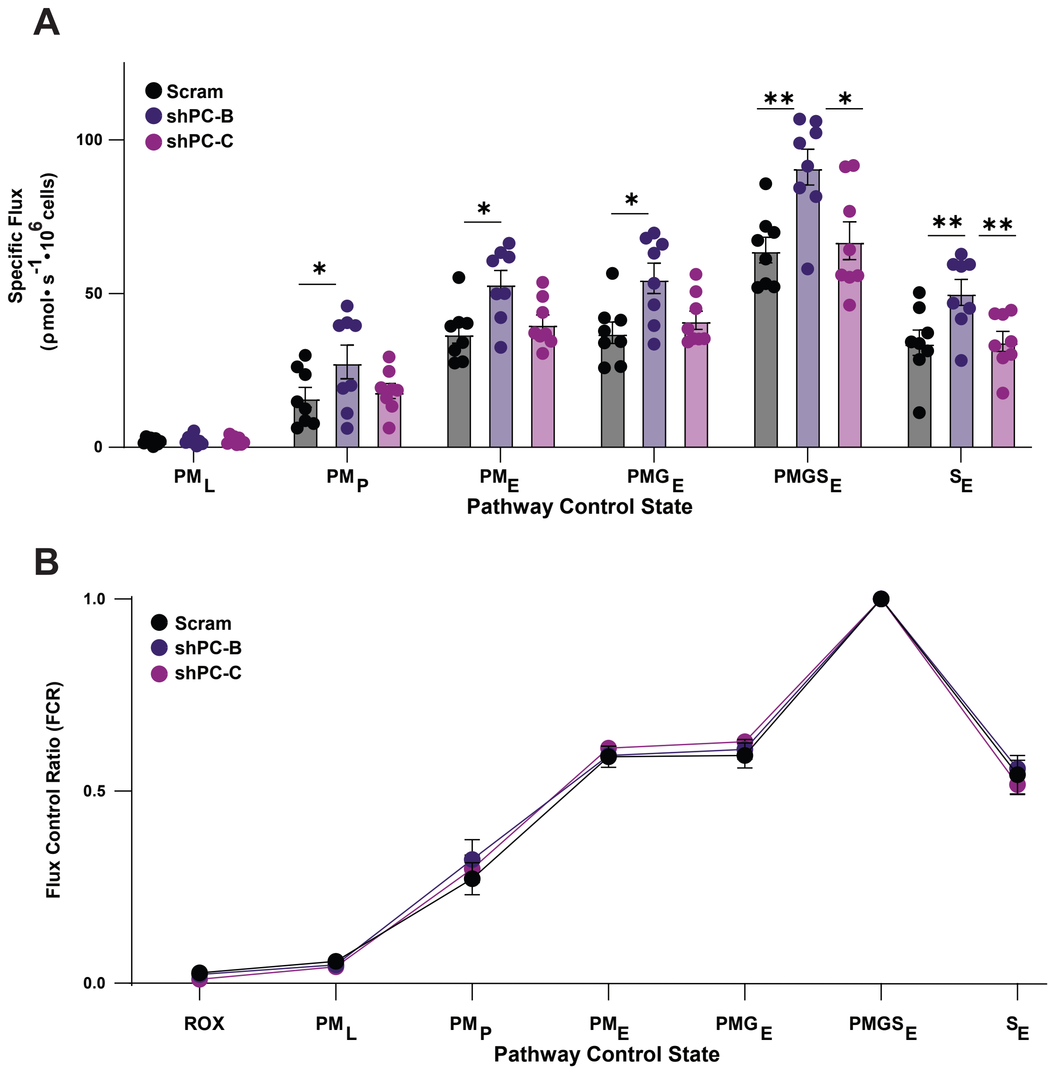
